# A unified hypothesis-free feature extraction framework for diverse epigenomic data

**DOI:** 10.1101/2023.01.26.525794

**Authors:** Ali Tuğrul Balcı, Maria Chikina

## Abstract

**Motivation:** Epigenetic assays using next-generation sequencing (NGS) have furthered our understanding of the functional genomic regions and the mechanisms of gene regulation. However, a single assay produces billions of data represented by nucleotide resolution signal tracks. The signal strength at a given nucleotide is subject to numerous sources of technical and biological noise and thus conveys limited information about the underlying biological state. In order to draw biological conclusions, data is typically summarized into higher order patterns. Numerous specialized algorithms for summarizing epigenetic signal have been proposed and include methods for peak calling or finding differentially methylated regions. A key unifying principle underlying these approaches is that they all leverage the strong prior that signal must be locally consistent.

**Results:** We propose *L*_0_ segmentation as a universal framework for extracting locally coherent signals for diverse epigenetic sources. *L*_0_ serves to both compress and smooth the input signal by approximating it as piece-wise constant. We implement a highly scalable *L*_0_ segmentation with additional loss functions designed for NGS epigenetic data types including Poisson loss for single tracks and binomial loss for methylation/coverage data. We show that the *L*_0_ segmentation approach retains the salient features of the data yet can identify subtle features, such as transcription end sites, missed by other analytic approaches.

**Availability:** Our approach is implemented as an R package “l01segmentation” with a C++ backend. Available at https://github.com/boooooogey/l01segmentation.

## Introduction

Using next-generation sequencing (NGS) to profile the epigenetic states of biological systems has rapidly gained popularity. Large-scale efforts to profile the epigenome, such as ENCODE (3) and Roadmap (13), have paved the way for these assays to be used routinely to investigate diverse biological problems, from cancer biology to immune cell development.

Once the data is aligned, epigenetic assays can be represented as (possibly normalized) read densities along the chromosome. Without further processing, each experiment contains on the order of 3 *×* 10^9^ values but many downstream analyses make use of representations with a much smaller footprint. Many epigenetic assays produce signals that have peak-like structure, that is, sequencing reads pile up in specific chromosomal locations representing a small fraction of the genome. Such assays can be succinctly represented by recording the position and value of the peak. The MACS (25) peak discovery software is widely used by consortium projects. However, some histone marks, such as K27me3 and K27me36, do not conform to the peak-like structure but instead cover broad regions at the scale of genes. In these cases the approach taken by MACS is to heuristically merge narrow peaks (16) forming so-called “broad peaks”. Importantly, some assays cannot be described as either broad or narrow peaks and exhibit characteristics of both. For example, Pol2 signal is composed of a large peak at the transcription start site (corresponding the the paused polymerase state) and a “broad” peak covering the entire transcribed region (corresponding to active transcription).

Sequence-based methylation likewise does not conform to the assumption of peak-like structure. The methylation signal is locally coherent in the sense that the measurement of adjacent CpGs is correlated. This is likely to be governed by a common regulatory process, such as the binding of a TF or polymerase or gene transcription. However, the length of regions with consistent methylation is highly variable. Moreover, different patterns of local correlation structure emerges at different scales. While promoters of expressed genes are hypo-methylated they may still contain stretches of highly methylated DNA (see Figure 6 for an example). For experimental designs that are focused on specific biological contrasts, this local coherence of methylation is exploited by combining differentially methylated sites (DMSs) into stretches of differentially methylated regions (DMRs) using various heuristics (6; 19; 24; 1; 17).

Altogether, the key property of epigenetic data is that while the raw data is reported at nucleotide resolution nearby chromosomal positions are likely to be produced by the same biological or experimental process and this fact can be exploited to define narrow peaks, broad peaks, or DMRs. While these analysis approaches are popular and effective, they rely on specific assumptions, tunable parameters, or, in the case of DMRs, on knowledge of experimental design. In this work we propose a general-purpose framework that uses efficient *L*_0_ segmentation and assay specific loss functions to represent any NGS epigentic experiment as piece-wise constant while making no a priori assumption regarding the expected patterns in the data.

**Table 1.** Comparison of existing segmentation methods.

|  | Implementation Availability | Probability Distribution | Data Format Supported | Method class |
| --- | --- | --- | --- | --- |
| CROCS (14) | Yes | Gaussian, Poisson, Negative Binomial | ChIP-seq | Breakpoint detection with structured constraints (originally proposed as gfpop(18)). |
| PELT (12) | Yes | Flexible | Flexible | Break-point detection with pruned dynamical programming |
| Yokoyama et al (23) | No | Binomial | WGBS | Application of PELT |
| Herrmann et al (8) | No | Binomial ( signal vs. input) | ChIP-seq | Bayesian MCMC |
| Chen et al (2) | No | Gaussian | ChIP-seq | Bayesian MCMC |
| Xing et al (22) | No | Poisson | ChIP-seq | Hidden-markov model |

## Our Approach

A standard way of enforcing that signals are locally consistent is by penalizing the difference in adjacent values. Given an input vector *y*_*i*_ we seek to approximate it with a locally smoothed vector ***β*** by solving an optimization problem of the following general type:

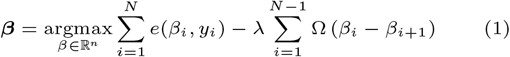

Here *e*(*β, y*) is a reconstruction error term, and Ω(*β*_*i*_ *− β*_*i*+1_) is a regularization term that penalizes the differences between consecutive values.

If we take squared error and the penalty to be the *L*_1_ norm this problem is known as fused-Lasso. This penalty drives the difference between successive elements of *β* to 0 and hence the approximation solution is piece-wise constant. Because of this property, the fused-Lasso can also be seen as a change-point detection algorithm.

In this work we consider a different penalty function, the *L*_0_ error. Instead of penalizing the sum of the absolute values of the differences, this simply penalizes the total number of change points regardless of their values. The *L*_0_ formulation has important performance and algorithmic consequences. Because all change points are penalized equally, the *L*_0_ formulation can produce much sharper breaks, which, as we show, allows for more efficient segmented representations. Another attractive property of the *L*_0_ formulation is that for the optimal solution, the value of the fused segment is purely a function of the individual data points in the segment (in the case of squared and Poisson error, it is the mean) and does not depend on the adjacent segments.

In terms of algorithmic considerations, unlike the *L*_1_ formulation, the *L*_0_ formulation is not convex. However, recent algorithmic advances allow for the solution to be found in empirical linear time (11).

Our implementation, which extends the approach described for solving *L*_0_ segmentation (11), uses error functions derived from different probability distributions. We implement the standard Gaussian distribution (squared error) as well as distributions that more accurately model epigenetic count data, such as Poisson for single-track data and binomial for double-track data counts (i. e. methylation and coverage). We find empirically that the *L*_0_ segmentation scales linearly when applied to biological data. In Fig. 1, we plot the time it takes to solve the *L*_0_ segmentation problem with different probability distributions for various input sizes.

**Fig. 1.**
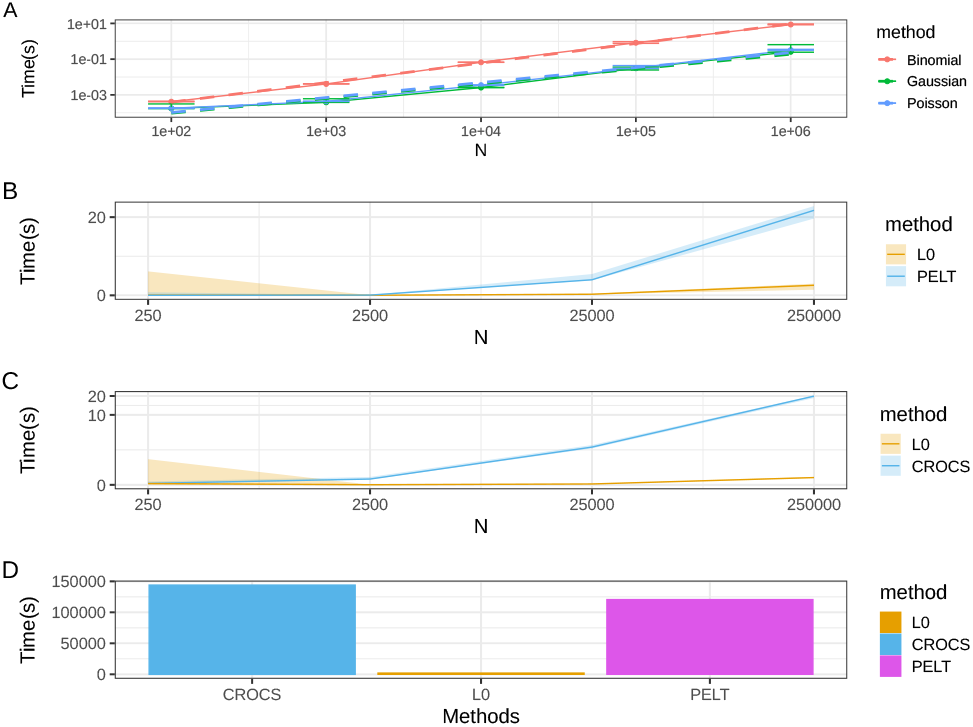
**A:** Binomial, Gaussian, and Poisson *L*_0_ segmentation execution times are plotted in log-space (both x and y) with respect to various segment lengths. The data is simulated using the corresponding probability distributions. Even though the theoretical worst case complexity of the algorithm is *O*(*n*^2^), in practice, the execution time scales linearly with the length of the input. **B** and **C:** comparison of the running time of our proposed method against similar approaches. CROCS is used for epigenetic data. PELT is the algorithm used in the Yokoyama et al. approach for segmenting methylation data(?) **D:** To estimate how long it would take for the methods to segment the entire Hg38 we benchmarked the methods on a 300KB segment and multiplied the results with 10000. This method highly likely underestimates both CROCS’ and PELT’s actual execution time.

**Fig. 2.**
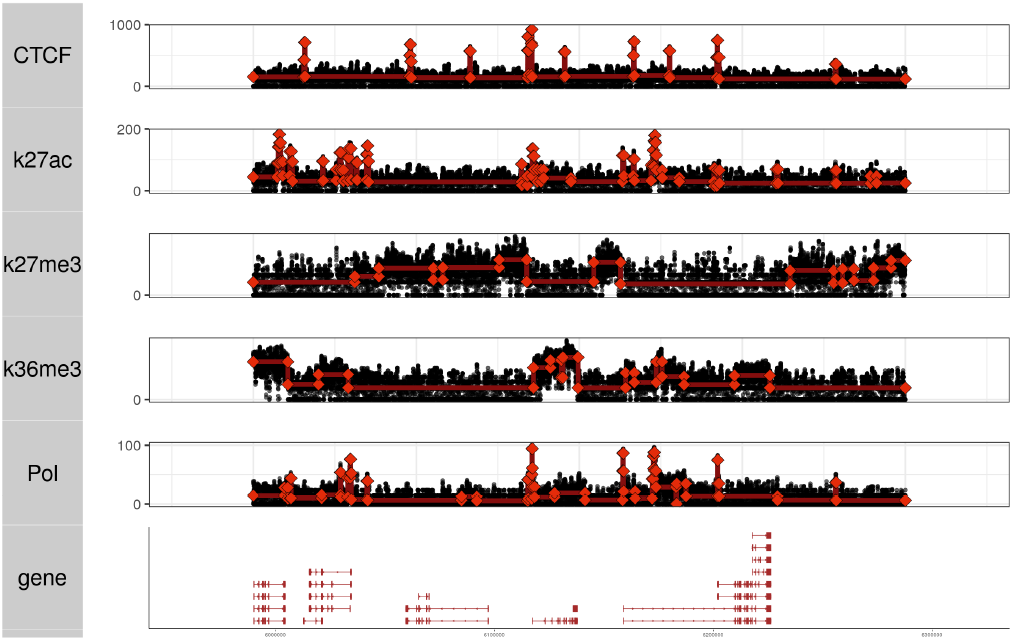
Poisson *L*_0_ segmentation on various epigenetic tracks is shown. The hyperparameter (*λ*) is chosen automatically using cross validation.

The empirical performance of our implementation both scales linearly and is overall very efficient, taking just 0.1 and 1 second (for Poisson and Binomial respectively) on 10^6^ input. The scalability makes it possible to compress epigentic data genome-wide. For example, reading in the entire nucleotide-level Chip-seq signal of Pol II from the BigWig file, collating it into 20 base pair bins, applying *L*_0_ segmentation, and storing the segmentation in a BedGraph file takes ∼90 seconds with 4 cores. The increased time complexity of binomial segmentations is mediated by the fact that most methylation analysis is done on CpGs, and there are only ∼30 million CpGs in the human genome.

### Relationship with prior work

A number of unsupervised approaches for segmenting epigenetic data have been proposed. These cover a diverse range of modeling assumptions and algorithmic approaches. Our work builds on these efforts in several important ways. Some previous methods have relied on Bayesian models using MCMC for inference. While this approach allows modeling flexibility, it is likely too time-consuming to be applied genome-wide. We note that the previously proposed Bayesian methods have no usable implementations, itself a major limitation, we infer that they are computationally intensive due to their reliance on MCMC sampling.

Our approach, on the other hand, is based on penalizing breakpoints and is conceptually similar to PELT and CROCS. However, the algorithmic approach is distinct, resulting in a significant speedup (see Fig. 1 B,C). Indeed using these existing *L*_0_ approaches we estimate it would take over a day to featurize an entire epigenetic track (Fig. 1 D). Our estimate is based on linear extrapolation is underestimate as these methods do not exhibit linear scaling (Fig. 1 B,C). We not that as these methods solve a similar optimization problem but have extensive time requirements we benchmark them only on the most difficult bench-marking task: transcription end site (TES) discovery and find that our implementation slightly outperforms PELT and significantly outperforms CROCS(Fig. S1).

Furthermore, we implement several useful error functions and can segment both single-track and double-track (e.g. methylation) data. We also provide a simple R package interface which allows direct segmentation of BigWig files.

We also note that in the context of methylation data analysis, it is common to use segmentation-type approaches to combine differentially methylated sites into differentially methylated regions (19; 20; 24; 1; 17; 5). These approaches rely on test statistics generated at the site level and generally cannot be applied out-of-the-box to the unsupervised segmentation of genome-wide signal.

### _*L*0_ segmentation retains salient genomic features

To illustrate features of *L*_0_ segmentation, we plot segmentation results across tracks with diverse features that represent different signal classes: CTCF and K27ac for narrow peaks, K27me3 and K36me for broad peaks, and Pol II, which shows a complex signal pattern composed of high narrow peaks and low broad peaks. Each track was segmented with *L*_0_ Poisson loss with the regularization parameter determined by optimizing “offset” cross-validation, a specialized cross-validation procedure that accounts for nuisance autocorrelation that arises from ChIP-seq fragments (see Methods).

We find that while neither the *L*_0_ segmentation nor the cross-validation procedure is optimized for a particular signal type, our approach automatically fits the expected patterns to each data type, effectively finding narrow peaks and broad peaks.

To demonstrate these qualitative observations in a quantitative evaluation framework, we make use of raw and processed datasets from the NIH Roadmap Epigenomics Consortium. The database provides an organized compilation of high quality uniformly processed raw epigenetic data as well as output form standard analysis tools such as MACS or ChromHMM (25; 4). Using these well-established analysis tools as a standard give us the opportunity to evaluate if we can distill different raw epigenetic tracks into a concise summary using a single principled algorithm. We focus on two cell lines that had all the relevant data types available: fibroblast cells (IMR90) and embryonic stem cells (H1). The *L*_0_ segmentation provides a reduced representation of a signal under the assumption that it is locally constant. As such, it doesn’t explicitly optimize specific features. Alternative approaches to compressing epigenetic datasets are equal-width local binning or using specialized algorithms to explicitly detect specific features of interest such as “peaks”. The MACS software (25) is highly optimized for peak discovery, and we use the MACS peak calls as a silver standard in order to evaluate the performance of segmented representtions.

Specifically, we evaluate if reduced representations capture the peak-like features with high fidelity by formalizing a notion of representation efficiency as the trade of between representation size and signal capture (defined in multiple ways). We evaluate *L*_0_ segmentation (our proposed solution), the convex *L*_1_ segmentation, and equal-size binning (a baseline that is commonly used to reduce epigenetic tracks in a data-agnostic way). We report the representation size in terms of “compression ratio,” which is the input length divided by the number of constant segments in the output.

In order to quantify how well the reduced representations retain peak features (signal fidelity), we formulate two different evaluations. The first one is based on computing the median fold change under the MACS peak regions after data compression. At the extreme value of compression ratio =1 (no compression) this just corresponds to the median fold change for the raw data. As the compression ratio is increased this number will eventually decrease as peak regions are merged with adjacent regions. The results of this evaluation are plotted in Fig. 3 (A,B). We find that according to this metric the *L*_0_ method is able to maintain accuracy up to 10,000-fold compression while both the binning and L1 show considerable drop-offs in performance much earlier.

**Fig. 3.**
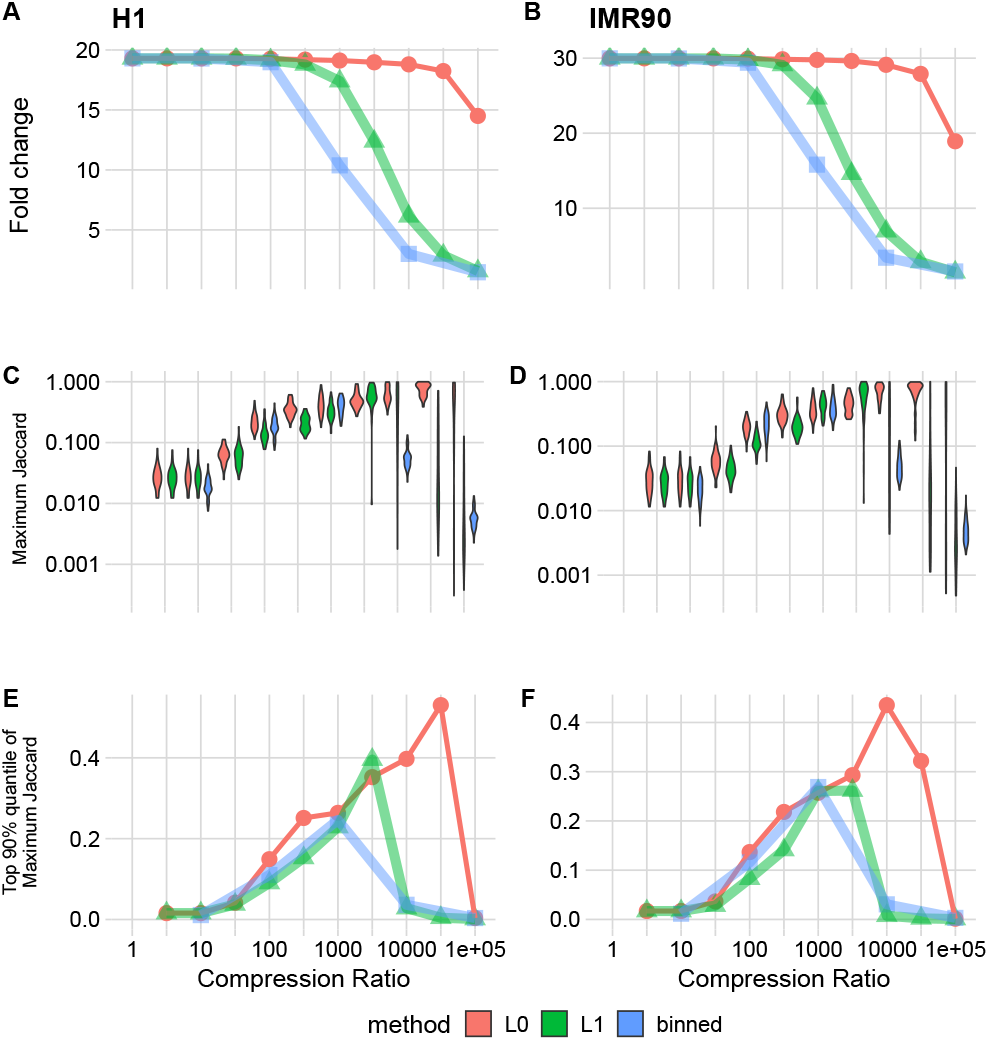
Comparison of *L*_0_Poisson (red), L1 Poisson (green), and fixed size binning (blue) with respect to the various compression ratios on the DNAse track from the H1 and IMR90 cell lines. The methods are applied on a 10M base-pair segment extracted from each cell line’s Chromosome 1. We evaluated the methods using 937 peaks for H1 and 1400 peaks for IMR90 discovered by MACS within this segment. **A and B:** The ratio of the mean signal within peak regions to the mean background signal after segmentation. **C and D:** Maximum Jaccard Index distributions for the methods at different compression ratios. Jaccard indices are calculated between peaks and the segments discovered by the methods. **E and F:** Median of the distributions shown in C and D.

The second evaluation is based on quantifying to what extent the reduced representation approximated the boundaries (as opposed to the values) of the peak calls. For each MACS peak we compute the maximum Jaccard index across all possible overlapping segment region. The distributions are plotted in Fig. 3 C,D and 90% quantile trends are plotted in the third row in panels E and F. In this evaluation, we see that the input (compression ratio =1), where each point corresponds to a segment is actually not optimal. This is expected as for the input each point is a single nucleotide and thus will not have a high Jaccard index with peak regions that are on the order of hundreds of nucleotides long. Instead, the optimal is reached at an intermediate value for all methods though with widely different compression ratios. Considering the H1 cell-line (C,E), local binning achieves optimal signal at a compression ratio of about 1000-fold, where 90% of peaks have a Jaccard concordance value of 0.2. For *L*_1_ the corresponding values are about 3000-fold and 0.4, a considerable improvement. On the other hand, *L*_0_ achieves both the best quantile value of 0.5 and does so at a substantially higher compression rate of 3 *×* 10^4^.

### *L*_0_ segmentation effectively captures complex patterns

In the previous section we evaluated segmented representations against MACS peak which serves to illustrate that the approach captures peak-like structure with high fidelity without making any assumptions. However, we do not expect to replace highly effective peak finding methods, rather we aim to show that we can discover salient features in datasets in an assumption free way. This in turn would enable us to analyze complex signals that are not easily summarized as peaks.

To do so, we focus on segmenting Pol II tracks. Pol II signals are composed of sharp peaks and the transcription start site and broad coverage along the transcribed gene length. As can be seen in the example shown in Fig. 4 B the segmentation procedure identifies both the sharp peak at the transcription start site and the broad transcript associated regions.

**Fig. 4.**
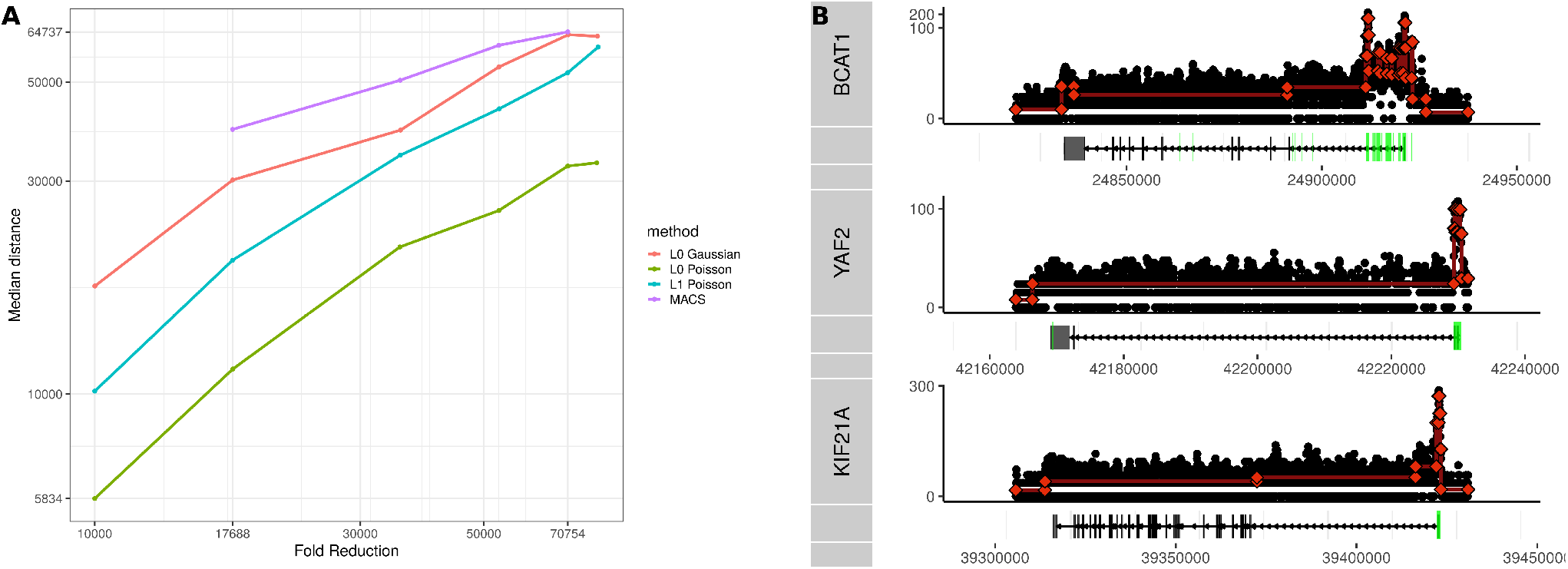
**A:** Comparison of the performance of different data reductions approaches on the transcript end site (TES) discovery task. We plot the median absolute distance to TES based on known gene models. The degree of segmentation is adjusted using lambda while for peaks we adjust the FDR cutoff to get a comparable number of segments. **B:** Examples of segmentation tracks for selected transcripts. Green regions indicate MACS peaks. While the Pol II signal is often noisy outside of the promoter region, a subtle drop-off in signal can often be observed towards the end of transcripts. In many cases this change in signal is correctly identified by L0 Poisson segmentation.

In particular, we expect that a correct segmentation of this signal should place a breakpoint near the transcription end site (TES). In Fig. 4 A, we quantify this via the median distance (MD) between a TES and the closest breakpoint using different data representations. We consider segmented data using either *L*_0_ or *L*_1_ penalty and Poisson or Gaussian distributions. The degree of compression is adjusted by the penalty parameter. We also consider the MACS peak based representation and vary the total number of peak based segments by adjusting the FDR cutoff. We demonstrate that the L0 Poisson model consistently achieves the lowest MD, with L0 Gaussian having the next best performance. In Fig. 4 B, we provide some examples of Pol II tracks and corresponding segmentations, demonstrating that *L*_0_ segmentation effectively finds TES-associated breakpoints which are missed by the MACS peak-based analysis (green). We also apply the TES discovery evaluation to the previous proposed break-point methods PELT and CROCS and find that despite solving a similar problem *L*_0_ slightly outperforms PELT. We note that CROCS does not perform well on this task as the additional constraints and the automatic hyper-parameter tuning are optimized for peak finding and not well suited for the data (Fig. S1).

### *L*_0_ compression for methylation data

Finally we consider the problem of featurizing DNA methylation. Most methods to extract higher order features from methylation data rely on grouping differentially methylated sites (DMSs) into differentially methylated regions(DMRs) post hoc (6; 19; 24; 1; 17). Thus, these need a specific statistical contrast to be specified and cannot be applied to a single track. Moreover, methylation data shows complex local correlation patterns that manifest at different scales and no standard approach (akin to MACS) to extract unsupervised features exists.

Unlike other epigenetic tracks, methylation is truly nucleotide resolution as it directly measures the methylation status of individual cytosines. Methylation data is reported as a count pair of methylated and unmethylated reads. The probability that a base is methylated (the ratio of methylated to total reads) is referred to as *β*. Importantly, the methylation signal doesn’t have natural “peaks” that are the hallmark of other epigenetic tracks such as ATAC-seq and ChIP-seq but is rather dispersed through the genome. Because the read coverage is scattered across the genome, the coverage at any single nucleotide is generally low but variable.

### Extending the L_0_ segmentation to binomial loss

NGS methylation data can be modeled as a series of draws from a binomial distribution with an unknown success probability. Considering the dual-track input, where 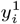 is the number of methylated reads and 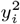 is the number of total reads, it can be derived as follows.

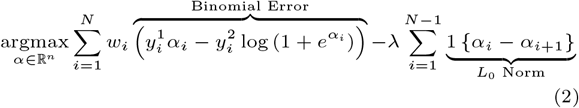

Explicitly, modeling the binomial error accurately accounts for read coverage at each location, so that low-coverage segments are more likely to be merged with their neighbors. Secondly, because the *L*_0_ does not depend on the size of the breakpoints, this formulation readily creates single CpG segments if the methylation level is different from that of their neighbors and coverage is sufficient. We demonstrate these two properties of our formulation on a simulated dataset in Fig. 5. In panel A, we simulate a region that has a dramatically different *β* from its neighborhood, but the value is not reliable because the coverage is low. Segmenting the *β* values directly using squared error assigns this segment its own region. On the other hand, using the explicit two-track binomial formulation, this region is merged with its neighbors (both segmentations have exactly three segments). In panel B, we simulated a constant methylation region with two single CpG outlier segments, which are readily discovered by *L*_0_ but not by *L*_1_/fused-lasso.

**Fig. 5.**
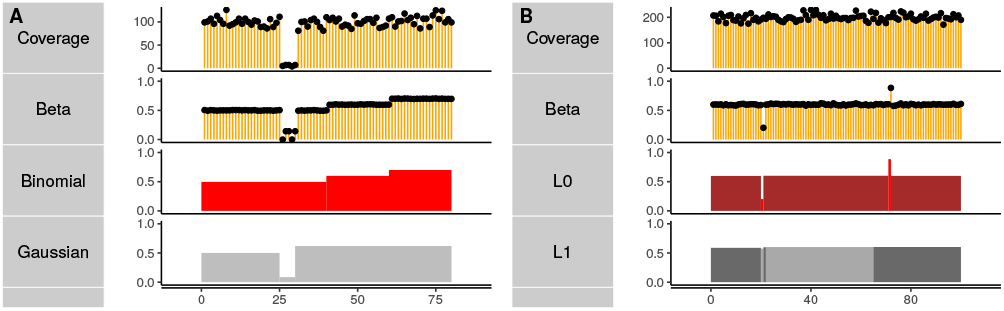
Binomial *L*_0_segmentation accurately accounts for read coverage and discovers short regions with distinct methylation rates. We compare segmentation formulations on simulated data. Methylation and coverage are shown in yellow. **A:** Squared (gray) and binomial error segmentation (red) with the same number of segments. Binomial error merges low coverage regions with their neighbors. **B:** Comparing *L*_0_ (red) and *L*_1_ (gray). The *L*_0_ penalty is more sensitive to local structure discovering the 2 CpGs simulated with different *β*.

**Fig. 6.**
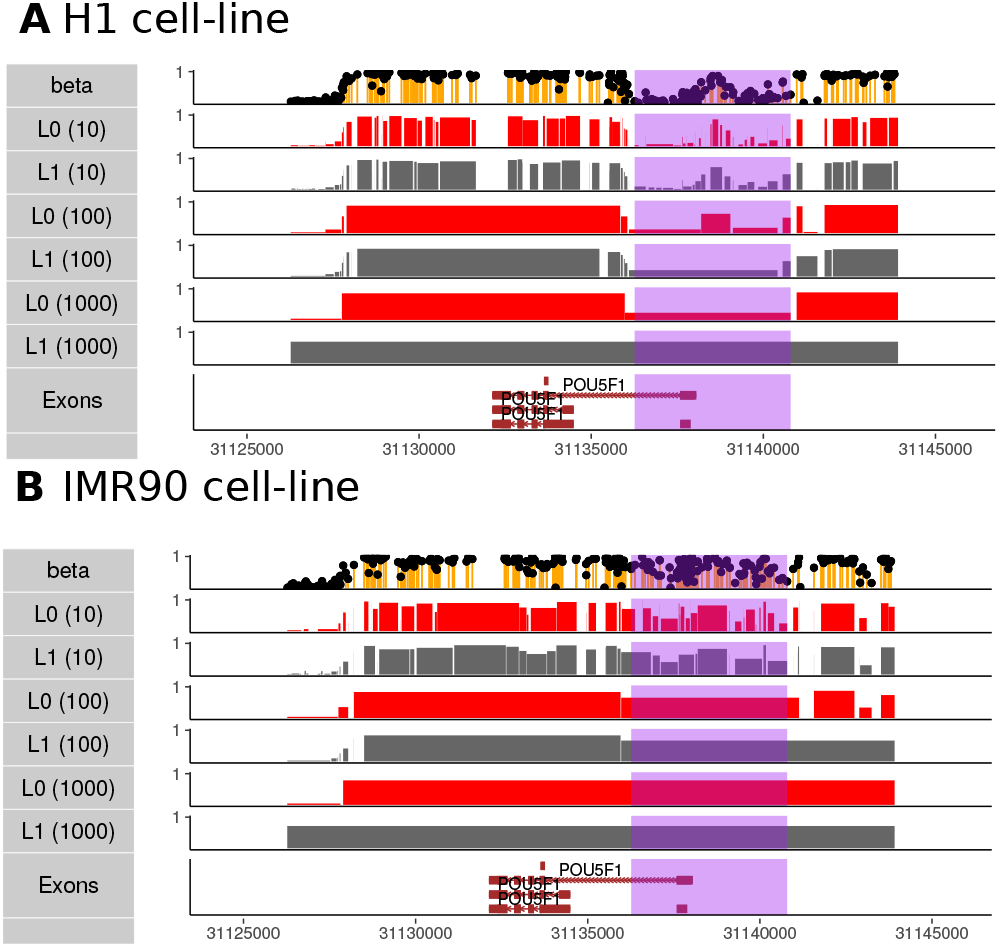
Comparing *L*_0_ to *L*_1_ with binomial error on real WGBS data with different fold compression computed on the chromosome level. The reported differentially methylated region is highlighted in purple. **A:** POU5F1 expressing cell line, H1. **B:** The non-expressing contrasting cell-line, IMR90. Note that unlike Fig. 5, fold compression is fixed at the chromosome level and consequently the number of segments in this region is not constant across *L*_0_ and *L*_1_.

We further illustrate the properties of binomial *L*_0_ segmentation on real WGBS data in H1 and IMR90 cell lines. We consider a region near the POU5F1 locus, which is an important pleuripotency gene and a major differentially methylated region between these two cell lines highlighted in the original WGBS study (15). In Fig. 6, we consider different segmentations of this region, fixing chromosome-level fold reduction, and we contrast *L*_0_ and *L*_1_. For reference, the differentially methylated region (DMR) of the POU5F1 promoter, discovered with supervised analysis, is shown in purple. Contrasting the *L*_0_ and *L*_1_ formulations, we see that the unsupervised *L*_0_ segmentation is able to identify the promoter DMR in the POU5F1-expressing cell line, H1, at up to 1000-fold compression. The promoter DMR is identified as a feature up to 100-fold reduction in the non-expressing cell line IMR90.

The region around POU5F1 shows a common methylation pattern of expressed genes, where the promoter tends to be hypomethylated while the gene body is hypermethylated. We thus evaluated how well these patterns are preserved globally using compressed representations. Similarly to our previous evaluations of compression for other epigenetic tracks, we evaluate the trade-off between compression level and signal retention, focusing on the chromoHMM states denoted as TssA (TSS) and Tx (transcript). As these are derived from other epigenetic tracks from the same cell line, they mark only the genes that are expressed in each. We find that using raw methylation input, we observe the expected hypomethylation for TssA and hypermethylation for Tx. The pattern is nearly unaltered as we increase the compression level up to 1000-fold for *L*_0_ (see Fig. 7). In contrast, for *L*_1_/fused-lasso, the TssA enrichment begins to decrease at a compression ratio of 100. In summary, we find that the binomial error *L*_0_ segmentation we propose is an effective unsupervised feature reduction methods for NGS methylation data. It naturally accounts for read coverage, can extract very small segments with distinct methylation patterns, and discover salient epigenetic features such as promoters and DMRs in a fully unsupervised manner.

**Fig. 7.**
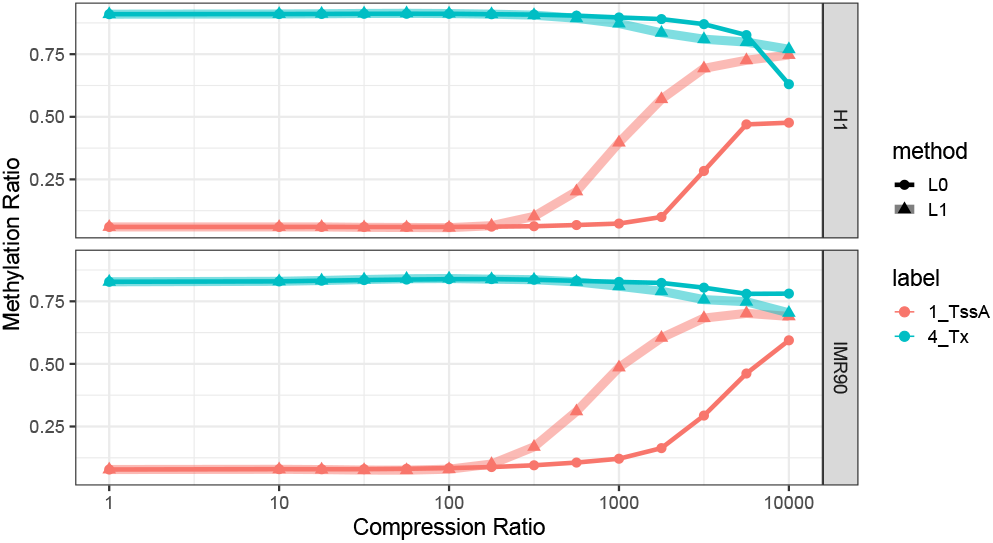
Comparison of *L*_0_ and *L*_1_ segmentations with binomial error in preserving the characteristics of genomic regions, namely “TssA” (TSS) and “Tx” (Transcript) regions identified by ChromHMM, in two cell lines: H1 and IMR90.

## Conclusion

Epigenetic data refers to a heterogeneous class of assay types that differ in their signal properties. However, all epigenetic data shares a local coherence structure as assay values at nearby nucleotides are determined by a common biological or technical process. This local structure can be leveraged in different ways via algorithms that are specialized for specific data types and data features.

Here, we propose a general computational approach that can exploit local structure in an entirely data-agnostic way. Our approach generalizes a recently proposed highly efficient *L*_0_ segmentation algorithm to probability distributions relevant to modeling epigentic data. We show that the *L*_0_ segmentation efficiently represents epigenetic tracks while retaining many salient features such as peaks, ChromHMM states, as well as transcription start and end sites making no assumptions about the underlying signal structure beyond piece-wise constant. This makes our approach suitable for representing diverse track types, including those that have “broad peak” or even complex structure such as Pol II, and methylation in a single unifying framework.

## Methods

### Data

All of the datasets used in our analysis are from the IMR90, H1, or K562 cell lines. The example dataset depicted in Fig. 2 is a subset of the K562 BigWigs used in the Segway (9) analysis and was downloaded from https://noble.gs.washington.edu/proj/segway/2011/bigWig/. We used the following tracks: “CTCF.Bernstein”, “H3K27ac”, “H3K27me3”, “H3K36me3”, “Pol2 8WG16.Snyder.” For our evaluations of ChIP-seq data, we used raw data, MACS peak calls, and ChromHMM results from the H1 and IMR90 cell lines. These datasets are available in the Roadmap data portal at https://egg2.wustl.edu/roadmap/data/byFileType/alignments/ for raw data, https://egg2.wustl.edu/roadmap/data/byFileType/peaks/ for MACS peaks, and https://egg2.wustl.edu/roadmap/data/byFileType/chromhmmSegmentations/ for ChromHMM segmentations. For WGBS, we used the fraction and coverage tracks available at https://egg2.wustl.edu/roadmap/data/byDataType/dnamethylation/WGBS/FractionalMethylation_bigwig/ and https://egg2.wustl.edu/roadmap/data/byDataType/dnamethylation/WGBS/ReadCoverage_bigwig/. For the evaluation of transcription end site (TES) discovery, we used the K562 used Pol II assay BigWig and BAM files provided by the ENCODE data portal with accession code ENCSR000BM.

### Software

*L*_0_ segmentation is implemented as an R package with a C++ backend. The embedding of C++ in R is carried out by the Rcpp interface. The implementation is available at https://github.com/boooooogey/l01segmentation. Within R, `fusedsegmentation` is the main function that delegates segmentation tasks to various solvers. Through the arguments, users can specify a distribution (Gaussian/Poisson/binomial), a penalty (*L*_0_/*L*_1_), and change the format of the output. Another valuable function is `compressBigwig`, an end-to-end pipeline, that takes a BigWig file as input and generates a BedGraph with segments. Detailed information about the algorithm and our implementation can be found in the appendix.

### Binned cross-validation

Cross-validation can be used to find optimal regularization parameters for fused lasso and related segmentation problems (21; 10). In the standard cross-validation procedure, individual points are left out of the computation and estimated using the value of their segment. The best *lambda* is taken to be the one that achieves minimal test error. If the data are independently sampled from the piece wise constant signal, this procedure will identify the correct *λ* that recovers the underlying structure without capturing the sampling noise. In the case of epigenetic assays that depend on immunoprecipitation (ChIP), independent sampling conditioned on true signal is not an accurate assumption. Since the protein is pulled down with a DNA fragment that is much larger than its actual occupancy site, regions of the DNA that are not directly bound but are simply near the site are likely to be captured. Moreover, the probability of capture depends continuously on the proximity to the true occupied site, inducing local dependency across adjacent sites and violating independent sampling.

Instead, the value at a single position is best predicted as the average of its immediate neighbors, and using naive cross-validation results in a very large number of segments. To avoid fitting this local structure, we propose a binned cross-validation strategy. Instead of leaving out random or equally spaced elements, the data is divided into windows whose width approximately matches the expected ChIP-seq fragment length (we use 300bp), and the test set is then taken to be equally spaced windows. Furthermore, the test region signal is reconstructed as the average of the left-most point in the adjacent left window and the right-most point in the adjacent right window.

To illustrates, consider a region of length 6 and a cross-validation window of 2. One of the cross-validation test-train splits will be as follows: train: *X*_train_ = *X*_1_, *X*_2_, *X*_5_, *X*_6_ and test: *X*_test_ = *X*_3_, *X*_4_. We perform segmentation on *X*_train_ and we predict 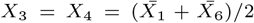, where 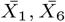 are the predicted values.

We note that for methylation data, independent sampling is valid and regular cross-validation produced intuitive results.

### Transcription end site (TES) analysis

For the transcription end site (TES) discovery task, we utilized the BAM and BigWig files for K562 Pol II signal obtained from ENCODE accession code ENCSR000BMR. These BAM files were merged using the `merge` function from `samtools` and subsequently converted into BigWig files using `bamCoverage`. The L0/1 segmentation framework was applied to the BigWig format, while MACS was employed on the BAM files.

To assess the impact of distinct objective functions and regularizers, we segmented the Pol II signal using the Gaussian and Poisson distributions along with the L0 regularizer, as well as the Poisson distribution with the L1 regularizer. To ensure comparable segmentations across different assumptions, we adjusted the hyperparameter *λ*. Since broad peaks were not available in the Encode portal we reran the MACS analysis as follows:

~~~
macs3 callpeak -t rep1_rep2_combined.bam -c
     ctrl1_ctrl2_combined.bam --broad -g hs --broad-cutoff 0.1
    -n broad_peaks_pol2
macs3 callpeak -t rep1_rep2_combined.bam -c
     ctrl1_ctrl2_combined.bam -f BAM -g hs -n test -B -q 0.01
~~~

Using mostly the default values. The broad and narrow peaks were merged for the evaluation. To enable a fair comparison between MACS peaks and segments, we treated the gaps between peaks as additional segments. During the evaluation, the peaks were sorted based on the adjusted p-values obtained from MACS, and an equivalent number of peaks were selected from the top for comparison.

The test was conducted on a 30 million base pair (bp) segment extracted from the hg38 reference genome, specifically Chromosome 12, starting from the genomic location 8 million bp. The RefSeq gene annotation dataset was utilized as the reference for identifying transcription end sites (TESs). Initially, 807 TESs were identified. Subsequently, additional filtering was performed based on the Pol II activity at the corresponding transcription start site (TSS) regions. As a result of these steps, 600 TESs remained.

MACS identified 848 peaks within the designated region, which were subsequently divided into four equal parts. In each evaluation step, an additional quarter of the peaks was included. To ensure an equal number of segments for each method, we adjusted the hyperparameter lambda for the L0/1 framework accordingly. The nearest function from the GenomicRanges library was employed to calculate the nearest breakpoints for each transcription end site (TES).

### Time Comparisons

The evaluation of L0 segmentation using Gaussian and Poisson distributions was performed on the raw read counts derived from BAM files of the same ENCODE experiment as described earlier (accession code ENCSR000BMR). However, for L0 segmentation using the binomial distribution, the evaluation was conducted on the methylation signal obtained from whole-genome bisulfite sequencing (WGBS) data extracted from adult mouse heart tissue, which was part of the dataset published by He et al (7).

Both the methylation and Pol II signals were analyzed using sections of varying lengths. The sections ranged from 100 base pairs (bp) up to 1 million bp, with each step increasing the length by a factor of 10. These sections were extracted from Chromosome 10, starting from the 10 million bp position.

Throughout the experiment, a hyperparameter lambda of 50 was arbitrarily set. It’s worth noting that the value of this hyperparameter does not impact the runtime of the framework.

To compare the CROCS, PELT, and L0 frameworks, we utilized the same ENCODE experiment focusing on POL II. Our tests were conducted on Chromosome 1, using sections of varying lengths. The sections started at 250 base pairs (bp) and ended at 250,000 bp, with each step increasing the length by a factor of 10.

For CROCS, we downloaded the implementation from the GitHub repository `aLiehrmann/CROCS` and used the default parameter values during the experiments.

Regarding PELT, we did not have access to the exact implementation used by Yokoyama et al (23). However, we were able to find an implementation in the `changepoint` library. We set the penalty type to `“Manual”`. Increasing the penalty hyperparameter leads to a more extensive pruning process, which results in longer execution times. Considering our focus on biologically meaningful regions, we determined that a penalty parameter value of 50000 produced meaningful peaks for POL II in a small example. Consequently we decided to set the penalty parameter of PELT to this value throughout the time comparison study.

To ensure comparability between the L0 segmentation results and PELT, we set the hyperparameter lambda for L0 in such a way that it resulted in the same number of breakpoints as obtained by PELT.

## Author contributions statement

M.C. and T.B. conceived the approach, developed algorithms, analyzed the results, and wrote the paper.

---

**Fig. S1.**
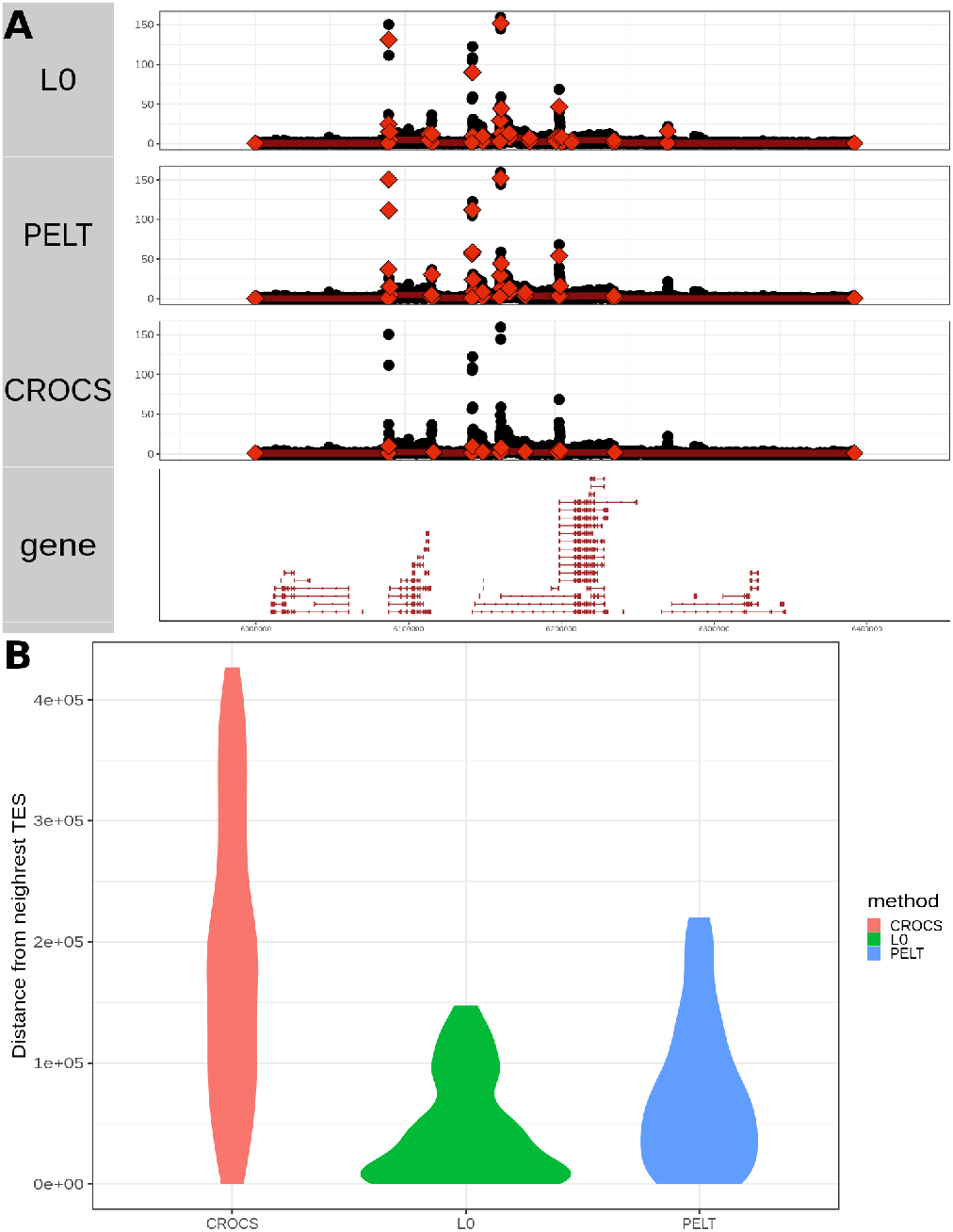
**A**: Illustration of segmentations of RNA Pol II using three different methods. CROCS, PELT, *L*_0_ have identified 13, 24 and 24 segments respectively. **B**: Distribution of distances from 148 transcription end sites identified within the 3Mb long genomic region starting from eight millionth nucleotide in chromosome 12 of Hg38 to the nearest breakpoints identified by one of the methods from a RNA Pol II assay. The hyper parameters for PELT and *L*_0_ are adjusted so that both algorithms produce the same number of break points; for the figure both algorithm produced 122 breakpoints. CROCS adjusts hyper parameters itself, and outputted three different segmentations for this segment. We evaluated the finest segmentation which has 9 breakpoints.

